# Conservation and Expansion of Transcriptional Factor Repertoire in the *Fusarium oxysporum* Species Complex

**DOI:** 10.1101/2023.02.09.527873

**Authors:** Houlin Yu, He Yang, Sajeet Haridas, Richard D. Hayes, Hunter Lynch, Sawyer Andersen, Gengtan Li, Domingo Martínez-Soto, Shira Milo-Cochavi, Dilay Hazal Ayhan, Yong Zhang, Igor V. Grigoriev, Li-Jun Ma

## Abstract

The *Fusarium oxysporum* species complex (FOSC) includes both plant and human pathogens that cause devastating plant vascular wilt diseases and threaten public health. Each *F. oxysporum* genome comprises core chromosomes (CCs) for housekeeping functions and accessory chromosomes (ACs) that contribute to host-specific adaptation. This study inspected global transcription factor profiles (TFomes) and their potential roles in coordinating CCs and ACs functions to accomplish host-specific pathogenicity. Remarkably, we found a clear positive correlation between the sizes of TFome and proteome of an organism, and FOSC TFomes are larger due to the acquisition of ACs. Among a total of 48 classified TF families, 14 families involved in transcription/translation regulations and cell cycle controls are highly conserved. Among 30 FOSC expanded families, Zn2-C6 and Znf_C2H2 are most significantly expanded to 671 and 167 genes per family, including well-characterized homologs of Ftf1 (Zn2-C6) and PacC (Znf_C2H2) involved in host-specific interactions. Manual curation of characterized TFs increased the TFome repertoires by 3%, including a disordered protein Ren1. Expression profiles revealed a steady expression of conserved TF families and specific activation of AC TFs. Functional characterization of these TFs could enhance our understanding of transcriptional regulation involved in FOSC cross-kingdom interactions, disentangle species-specific adaptation, and identify targets to combat diverse diseases caused by this group of fungal pathogens.

## INTRODUCTION

The fungal species complex of *Fusarium oxysporum* (FOSC) has been used as a model to study cross-kingdom fungal pathogenesis. Members within FOSC can cause devastating fusarium wilt diseases among economically important crops (Ma et al. 2013; Ma 2014; Michielse and Rep 2009; Ploetz 2015; Edel-Hermann and Lecomte 2019; Pegg et al. 2019; Yang et al. 2020; Dean et al. 2012; Rahman et al. 2021; Viljoen et al. 2020; Halpern et al. 2018) and is listed among the top five most important plant pathogens that have a direct impact on the global economy and food security (Dean et al. 2012). With strong host specificity, plant pathogenic *F. oxysporum* strains are further grouped as *formae speciales* (Armstrong and Armstrong 1981). For instance, tomato pathogens are named *F. oxysporum* f.sp. *lycopersici*, cotton pathogens *F. oxysporum* f.sp. *vasinfectum* (Halpern et al. 2018), and banana pathogen *F. oxysporum* f.sp. *cubense* (Viljoen et al. 2020). Recently, members within FOSC have also been reported to be responsible for fusariosis, the top emerging opportunistic mycosis (Ma et al. 2013; Yang et al. 2020), and fusarium keratitis, one of the major causes of cornea infections in the developing world and the leading cause of blindness among fungal keratitis patients (Kredics et al. 2015; Hassan et al. 2016).

Comparative genomics studies on this cross-kingdom pathogen revealed that the FOSC genomes, both human and plant pathogens, are compartmented into two components: the core chromosomes (CCs) and accessory chromosomes (ACs). While CCs are conserved and vertically inherited to execute essential housekeeping functions, horizontally transmitted ACs are lineage- or strain-specific and related to fungal adaptation and pathogenicity, conferred by virulent factors such as SIX (Secreted in Xylem) proteins (Ma et al. 2013; Yang et al. 2020; Rep et al. 2004; Yu et al. 2023). ACs and CCs must coordinate their gene expression to coexist within the same genome.

A few characterized transcription factors (TFs) coordinate the crosstalk between CCs and ACs, two compartments. One intriguing example is the cross-regulation among *F. oxysporum* transcription factors Sge1 (SIX Gene Expression 1), Ftfs, and effector genes. Sge1 is a highly conserved, CC-encoding TF. By name definition, Sge1 regulates the expression of SIX proteins (Michielse et al. 2009; van der Does et al. 2016). AC-encoding Ftf1 proteins (Ftf1 and its AC homologs) and a CC-encoding Ftf2 (Ftf1 CC homolog) are reported in the reference genome of *F. oxysporum* f.sp. *lycopersici Fol*4287 (van der Does et al. 2016). Constitutive expression of either *Ftf1 genes* or *Ftf2* induced the expression of effector genes (van der Does et al. 2016). Furthermore, It was documented that DNA binding sites of Sge1 and Ftf1 are enriched among the cis-regulatory elements of *in planta* transcriptionally up-regulated genes (van der Does et al. 2016). Another CCs and ACs cross-talking example is the alkaline pH-responsive transcription factor PacC/Rim1p reported in *F. oxysporum* clinical strains (Zhang et al. 2020). In addition to the full-length *PacC* ortholog (*PacC_O*), located on a CC, the clinical isolate NRRL32931 genome encodes three truncated *PacC* homologs, named *PacC_a*, PacC_b, and *PacC_c* in ACs (Zhang et al. 2020).

To thoroughly understand the coordination of the crosstalk between genome compartments and their contribution to the cross-kingdom fungal pathogenesis, this study compared the repertoire of TFs (*i.e*., TFome) among 15 *F. oxysporum* and 15 other ascomycete fungal genomes, which was organized into 48 families based on the InterPro classification of proteins. Remarkably, we discovered a strong positive correlation (*y* = 0.07264*x* − 190.9, *r*^2^= 0.9361) between the number of genes (*x*) and TFome size (*y*) of an organism. Primarily due to the acquisition of ACs, we observed increased TFome sizes among FOSC genomes. Fourteen out of 48 families involved in transcription/translation regulations and cell cycle controls are highly conserved. Thirty, accounting for ¾ of all families, are expanded in various degrees among FOSC genomes. Unique TF expansions driven by ACs include members of Zn2-C6 fungal-type (Zn2-C6) and Zinc Finger C2H2 (Znf_C2H2) families. This comparative study first highlights the conserved regulatory mechanisms of *F. oxysporum*, which are essential for variability and plant colonization. With the foundation established by functional conservation, this study further emphasizes potential modifications of existing regulatory pathways by acquiring additional TFs. In combination with existing expression data, this study may provide clues to the fine-tuning of networks in the environmental adaptation of this group of diverse organisms to engage in complex cross-kingdom interactions with different hosts.

## MATERIALS AND METHODS

### Generation of fungal TFomes

The annotation pipeline is briefly summarized in Figure S1A-B. The fungal proteomes of 30 strains were downloaded from the JGI MycoCosm portal (Grigoriev et al. 2014). Protein annotation was performed using InterProScan/5.38-76.0 (https://www.ebi.ac.uk/interpro/search/sequence/) (Jones et al. 2014). Annotations of proteins putatively serve as TFs were filtered out using a table containing InterPro terms related to transcriptional regulatory functions summarized by literature (Park et al. 2008; Shelest 2017), with further addition by manual curation (Table S1). Orthologous analysis to probe orthologs of functionally validated TFs (Table S3-4 and Table 3) in *Fusarium* was done with OrthoFinder 2.5.4 (https://github.com/davidemms/OrthoFinder) (Emms and Kelly 2019).

### RNA-seq analysis

The RNA-seq datasets were previously described (Guo et al. 2021; Redkar et al. 2022) and deposited by those authors to the NCBI Short Read Archive with accession number GSE87352 and to the ArrayExpress database at EMBL-EBI (www.ebi.ac.uk/arrayexpress) under accession number E-MTAB-10597, respectively. For data reprocessing, reads were mapped to reference genomes of Arabidopsis [annotation version Araport11 (Cheng et al. 2017)], Fo5176 (Fokkens et al. 2021), Fo47 (Wang et al. 2020) and Fol4287 (Ma et al. 2010) using HISAT2 version 2.0.5 (Kim et al. 2019). Mapped reads were used to quantify the transcriptome by StringTie version 1.3.4 (Pertea et al. 2015), at which step TPM (transcript per million) normalization was applied. Normalized read counts were first averaged per condition and then transformed by log2 (normalized read count + 1) and Z-scaled, then visualized in pheatmap (version 1.0.12).

### Genome partition

The genome partition results for chromosome-level assemblies were retrieved from previous reports for Fol4287 (Ma et al. 2010), FoII5 (Zhang 2019), Fo5176 (Fokkens et al. 2021), and Fo47 (Wang et al. 2020). Fo47 has a clear genome partition with 11 core chromosomes and one accessory chromosome, therefore serving as the reference for the genome partition of other *F. oxysporum* genomes. mummer/3.22 was applied to align scaffolds of genome assemblies against 11 core chromosomes of the reference genome Fo47 using default parameters. The scaffolds aligned to the core chromosomes of Fo47 with a coverage larger than 5% were annotated as core scaffolds. The rest of the scaffolds were partitioned as accessory scaffolds. Genes residing on core and accessory scaffolds were annotated as core and accessory genes, respectively.

### Phylogenetics analysis

Protein sequences were aligned via MAFFT/7.313 (Katoh and Standley 2013). Then the iqtree/1.6.3 (Minh et al. 2020; Nguyen et al. 2015) was run on the sequence alignment to generate the phylogeny (by maximum likelihood method and bootstrapped using 1000 replicates) (Hoang et al. 2018) and was then visualized via the Interactive Tree of Life (Letunic and Bork 2021), producing the phylogram. OrthoFinder 2.5.4 (Emms and Kelly 2019) was used for orthogroup determination. To build a species phylogram, randomly selected 500 conserved proteins (single-copy orthologs) were aligned first. Then the alignment was concatenated, and phylogeny was determined and visualized using the above methods.

## RESULTS

### 1. FOSC TFome expansion resulted from the acquisition of ACs

We compared 30 ascomycete fungal genomes (Figure 1 and Table 1), including 15 strains within the FOSC, nine sister species close to *F. oxysporum*, two yeast genomes (*Saccharomyces cerevisiae* and *Schizosaccharomyces pombe*), four other filamentous fungal species (*Neurospora crassa*, *Aspergillus nidulans, Aspergillus acristatulus*, and *Magnaporthe oryzae*). To maintain consistency, the protein sequences for all these genomes were retrieved from the MycoCosm portal (Grigoriev et al. 2014).

**Figure 1.**
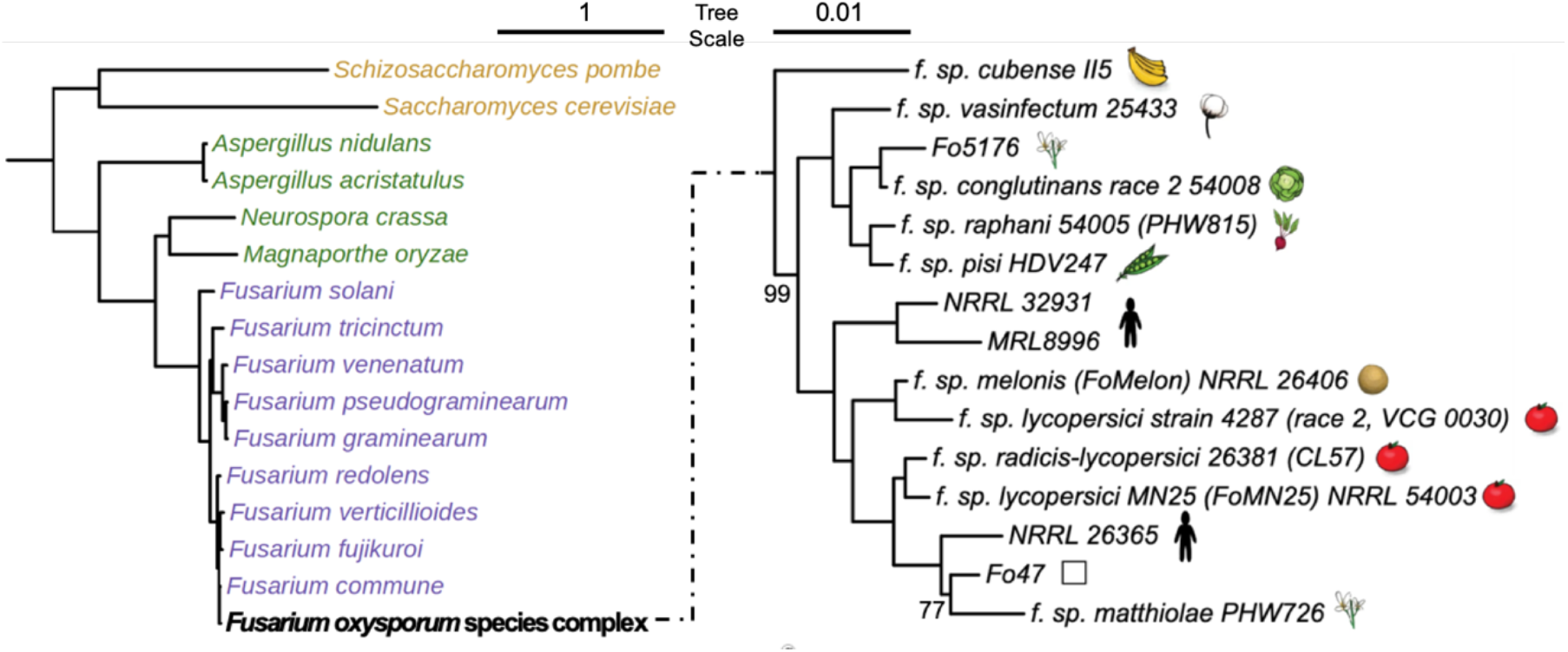
Phylogeny of fungal genomes included in this study. Both left and right phylograms were constructed by concatenated alignment of randomly selected 500 single-copy orthologous proteins, followed by the maximum likelihood method with 1000 bootstraps. Left shows a phylogram of FOSC (represented by the reference genome Fol4287) together with the other 15 ascomycetes. The right shows a phylogram of members within FOSC, rooted by *F. verticillioides* (not shown). Only bootstrap values not equal to 100 are shown.

**Table 1.** Fungal genomes used in this study.

| Fungal species or strains | MycoCosm identifier | Genome Size (MB) | No. of genes | TFome Size | Host | Reference |
| --- | --- | --- | --- | --- | --- | --- |
| *Saccharomyces cerevisiae* | Sacce1 | 12.07 | 6575 | 284 |  | (Goffeau et al. 1996) |
| *Schizosaccharomyces pombe* | Schpo1 | 12.61 | 5134 | 228 |  | (Wood et al. 2002) |
| *Aspergillus nidulans* | Aspnid1 | 30.48 | 10680 | 635 |  | (Galagan et al. 2005) |
| *Aspergillus acristatus* | Aspacri1 | 32.59 | 11221 | 666 |  | (Vesth et al. 2018) |
| *Neurospora crassa* | Neucr2 | 41.04 | 9730 | 447 |  | (Galagan et al. 2003) |
| *Magnaporthe oryzae* | Magor1 | 40.49 | 12673 | 520 | Rice | (Dean et al. 2005) |
| *Fusarium solani* | Fusso1 | 52.93 | 17656 | 1137 | broad hosts | (Mesny et al. 2021) |
| *F. pseudograminearum* | Fusps1 | 36.33 | 12395 | 627 | Wheat | (Gardiner et al. 2012) |
| *F. graminearum* | Fusgr1 | 36.45 | 13321 | 608 | Wheat | (Cuomo et al. 2007) |
| *F. venenatum* | Fusven1 | 37.45 | 12845 | 802 |  | (Mesny et al. 2021) |
| *F. tricinctum* | Fustr1 | 43.69 | 14106 | 925 | Broad hosts | (Mesny et al. 2021) |
| *F. verticillioides* | Fusve2 | 41.78 | 15869 | 917 | Corn | (Ma et al. 2010) |
| *F. fujikuroi* | Fusfu1 | 43.83 | 14813 | 901 | Broad hosts | (Wiemann et al. 2013) |
| *F. redolens* | Fusre1 | 52.56 | 17051 | 1098 | Broad hosts | (Mesny et al. 2021) |
| *F. commune* | Fusco1 | 48.37 | 15731 | 1012 | Broad hosts | (Mesny et al. 2021) |
| *F. oxysporum* f.sp. *cubense* (II5) | FoxII5 | 49.43 | 16048 | 1047 | Banana | (Zhang 2019) |
| *F. oxysporum* f. sp. *radicis-lycopersici* (CL57) | Fusoxrad1 | 49.36 | 18238 | 1151 | Tomato | (Delulio et al. 2018) |
| *F. oxysporum* Fo47 (Fo47) | FusoxFo47_2 | 50.36 | 16207 | 1082 |  | (Wang et al. 2020) |
| *F. oxysporum* f. sp. *lycopersici* (MN25) | Fusoxlyc1 | 48.64 | 17931 | 1119 | Tomato | (Delulio et al. 2018) |
| *F. oxysporum* NRRL26365 (NRRL26365) | Fox26365_1 | 48.46 | 16047 | 1036 | Human | (Yang 2020) |
| *F. oxysporum* f. sp. *melonis* (FoMelon) | Fusoxmel1 | 54.03 | 19661 | 1219 | Melon | (Ma et al. 2014) |
| *F. oxysporum* f. sp. *lycopersici* (Fol4287) | Fusox2 | 61.36 | 20925 | 1292 | Tomato | (Ma et al. 2010) |
| *F. oxysporum* NRRL32931 (NRRL32931) | Fusox32931 | 47.91 | 17280 | 1072 | Human | (Zhang et al. 2020) |
| *F. oxysporum* MRL8996 (MRL8996) | FoxMRL8996 | 50.07 | 16631 | 1057 | Human | (Zhang et al. 2020) |
| *F. oxysporum* f. sp. *matthiolae* (PHW726) | FoxPHW726_1 | 57.22 | 17996 | 1157 | Brassica | (Yu et al. 2020) |
| *F. oxysporum* f. sp. *vasinfectum* (FoCotton) | Fusoxvas1 | 52.91 | 19143 | 1189 | Cotton | (Delulio et al. 2018) |
| *F. oxysporum* f. sp. *pisi* (HDV247) | Fusoxpis1 | 55.19 | 19623 | 1229 | Pea | (Williams et al. 2016) |
| *F. oxysporum* f. sp. *raphani* (PHW815) | Fusoxrap1 | 53.5 | 19306 | 1132 | Brassica | (Delulio et al. 2018) |
| *F. oxysporum* f. sp. *conglutinans* (PHW808) | Fusoxcon1 | 53.58 | 19854 | 1142 | Brassica | (Delulio et al. 2018) |
| *F. oxysporum* Fo5176 (Fo5176) | FoxFo5176 | 67.98 | 19130 | 1236 | Arabidopsis | (Fokkens et al. 2021) |

To have a comprehensive TFome annotation, we started with reported InterProScan (IPR) terms associated with fungal transcriptional regulation (Park et al. 2008; Shelest 2017) and curated a mapping with updated IPR classification (interproscan version: 5.38-76.0) (Blum et al. 2021). In addition, we searched the IPR classification of protein families and obtained all other terms related to the transcriptional regulation activity. This resulted in 234 TF-related IPR terms (Table S1). Since most of the terms are initially defined in the mammalian systems, it was not surprising to see that overall, our fungal genomes are only associated with 71 IPR terms out of the total 234 TF-related IPR terms (Table S1, Materials and Methods, and Figure S1A-B for annotation pipeline). After filtering out 13 and 10 terms for redundancy (two terms describing the identical domain) and minimal presentation (< 4 among the 30 genomes), respectively, this comparative TFome study focused on the rest 48 IPR terms, which represented a total of 27967 TFs (Table S1-S2). Notably, 12 out of 48 terms were not reported to be affiliated with fungal transcriptional regulation by either Park et al. 2008 or Shelest 2017 (Table S1), adding values to our manual IPR term search.

Comparing the total number of genes in a genome (*x*) and the total number of TFs within that genome (*y*), we observed a strong positive correlation (*y* = 0.07264*x -* 190.9, *r*^2^= 0.9361) (Figure 2A). Among all genomes included in this study, FOSC TFomes are the largest, with an average of 1144 TFs per genome (Figure 2A, Table 1). After partitioning each FOSC genome into core and accessory regions (see Materials and Methods for details), we observed a positive correlation between the number of TFs encoded in the accessory chromosomal region of each strain (defined as accessory TFs hereafter) with the size of accessory genomes (Mb) (*y* = 17.239*x* + 3.553) (Figure S2), suggesting that accessory chromosomes contribute directly to the expanded TFome.

**Figure 2.**
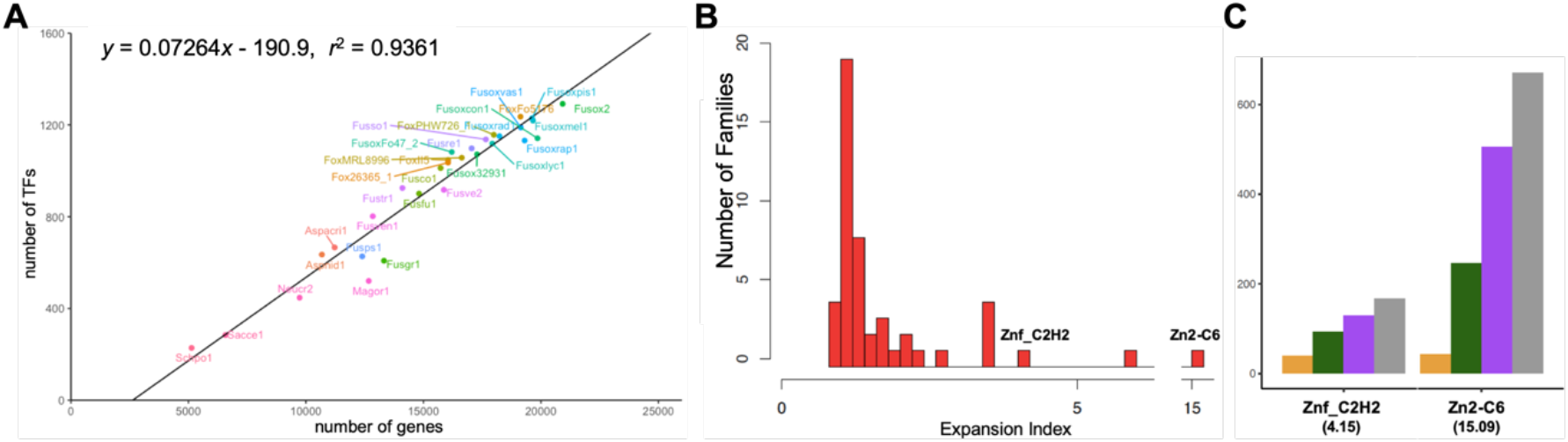
TFome conservation and variation among ascomycete fungi: baseline description. (A) There is a positive correlation between the number of genes and TFome size of an organism. JGI fungal genome identifiers were used as labels. (B) Histogram illustrates the distribution of expansion indexes among different families. (C) Average number of TFs of two most drastically expanded families (Znf_C2H2 and Zn2-C6) within each genome set. Genome Set 1 (G1) includes two yeast genomes (*S. cerevisiae* and *S. pombe*). Genome Set 2 (G2) includes four filamentous fungal species (*N. crassa*, *A. nidulans, A. acristatulus*, and *M. oryzae*). Genome Set 3 (G3) includes nine sister species close to *F. oxysporum*. Genome Set 4 (G4) includes 15 FOSC genomes.

To understand genome regulation among FOSC, we developed an expansion index score using two yeast lineages as the baseline (*EI_y_*):

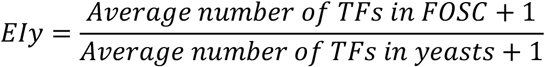

Based on this index value, we classified TF families into three major groups (Table 2, Table S1). Group 1 contains 14 TF families with an expansion score of 1, indicating high conservation. Group 2 includes four families with an index score of less than 1, reflecting some level of gene family contraction. Group 3 contains 30 families with an expansion index greater than 1, indicating gene expansion.

**Table 2.** Expansion Index (*EI_y_*) of 48 TF families. Asterisk indicates the families without a presence in yeasts

| IPR | Term | $El_y$ |
| --- | --- | --- |
| **Group 1** |  |  |
| IPR000814 | TBP | 1 |
| IPR003228 | TFIID_TAF12 | 1 |
| IPR004595 | TFIIH_C1-like | 1 |
| IPR006809 | TAFII28 | 1 |
| IPR042225 | Ncb2 | 1 |
| IPR008570 | Vps25 | 1 |
| IPR008895 | Vps72/YL1 | 1 |
| IPR007196 | CNOT1 | 1 |
| IPR005612 | CBF | 1 |
| IPR001289 | NFYA | 1 |
| IPR018004 | APSES-type HTH | 1 |
| IPR003150 | RFX | 1 |
| IPR033896 | MADS_MEF2-like | 1 |
| IPR018501 | DDT | 1 |
| **Group 2** |  |  |
| IPR006856 | MATalpha_HMGbox | 0.8 |
| IPR039515 | NOT4 | 0.9 |
| IPR033897 | MADS_SRF-like | 0.95 |
| IPR000232 | HSF | 0.98 |
| **Group 3** |  |  |
| IPR003163 | Tscript_reg_HTH_APSES-type | 1.04 |
| IPR001766 | Fork_head | 1.05 |
| IPR011016 | Znf_RING-CH | 1.11 |
| IPR001965 | Znf_PHD | 1.11 |
| IPR009071 | HMG_box | 1.12 |
| IPR004181 | Znf_MIZ | 1.24 |
| IPR001606 | ARID | 1.25 |
| IPR000679 | Znf_GATA | 1.3 |
| IPR001005 | SANT/Myb | 1.32 |
| IPR000818 | TEA/ATTS | 1.33 |
| IPR003120 | Ste12 | 1.33 |
| IPR003958 | CBFA_NFYB | 1.35 |
| IPR001083 | Cu_fist | 1.37 |
| IPR000967 | Znf_NFX1 | 1.4 |
| IPR006565 | Bromodomain | 1.52 |
| IPR001387 | Cro/C1-type_HTH | 1.6 |
| IPR001841 | Znf_RING | 1.64 |
| IPR000571 | Znf_CCCH | 1.74 |
| IPR001878 | Znf_CCHC | 1.83 |
| IPR010666 | Znf_GRF | 2 |
| IPR018060 | HTH_AraC* | 2 |
| IPR001356 | Homeobox | 2.28 |
| IPR007604 | CP2* | 2.73 |
| IPR007396 | PAI2 | 3.42 |
| IPR024061 | NDT80 | 3.47 |
| IPR011598 | bHLH | 3.48 |
| IPR007889 | HTH_Psq* | 3.53 |
| IPR013087 | Znf_C2H2 | 4.15 |
| IPR004827 | bZIP | 5.8 |
| IPR001138 | Zn2-C6 | 15.09 |

### 2. Conserved TF families that are primarily associated with general/global transcription factors

About 30% of the TF families, fourteen, are associated with strong orthologous conservation in all genomes we included in this study (Figure 2B; Table 2; Table S1). Because most of these conserved TF families are single-copied TF families, these 30% conserved TF families only account for less than 2% of the total TFomes. Based on a detailed study on *S. cerevisiae* and other model organisms, these TF families are involved in transcription/translation regulation and cell cycle controls.

#### 2.1. Transcription/Translation regulation

Either TF families are related to transcription initiation and elongation, including TATA box-binding protein (TBP), TBP-associated factors (TAFs), and RNA polymerase II elongation regulator Vps25. CCAAT-Binding Factors (CBFs) are related to ribosomal biogenesis. These families overall play conserved roles in general transcriptional and translational regulation across *Ascomycota*.

**Transcription initiation TBP** is one of the most conserved TF families, TBP binds directly to the TATA box to define the transcription start and initiate transcription facilitated by all three RNA polymerases. The function of TBP is so conserved as the yeast homolog can complement *TBP* mutations in humans (Yamaguchi et al. 2001; Roberts and Winston 1996).

**Transcription positive/negative regulators**, including **TAF12** and **TAF_II_28**, are parts of the transcription factor TF_II_D complex. Interacting with TBP, TAFs form the TF_II_D complex and positively participate in the assembly of the transcription preinitiation complex (Green 2000). Similarly, **TF_II_H** works synergistically with TF_II_D to promote the transcription (Fribourg et al. 2000). In contrast, Negative cofactor 2 (**Ncb2**) inhibits the preinitiation complex assembly (Goppelt et al. 1996). Other factors include the **CNOT1**, a global regulator involved in transcription initiation and RNA degradation (Chalabi Hagkarim and Grand 2020), and **Vps72/YL1** that contributes to transcriptional regulation through chromatin remodeling as reported in the yeast (Liang et al. 2016; Latrick et al. 2016).

**Transcription elongation: Vps25** is a subunit of the ESCRT-II complex, which binds to RNA polymerase II elongation factor to exert transcriptional control in mammalian systems (Kamura et al. 2001).

**Translational regulation**: CCAAT box is a common cis-acting element found in the promoter and enhancer regions of genes in the eukaryotes (Vuorio et al. 1990; Becker et al. 1991). **CBFs** are necessary for the 60S ribosomal subunit biogenesis and therefore involved in the translational control (Milkereit et al. 2001; Fromont-Racine et al. 2003; Edskes et al. 1998). This family, including Noc3, Noc4, and Mak21 in *S. cerevisiae*, has three members in each genome, and a clear single-copy orthologous relationship can be observed for each member (Figure S3A).

### 2.2. Cell cycle control

Five TF families are related to cell cycle control, including cell cycle progression, DNA repair, and machinery/cell integrity maintenance.

**APSES-type HTH** represents a family of fungal TFs involved in cell-cycle control and is crucial to the development (Xin et al. 2020). Every genome maintains four copies of genes encoding APSES-type HTH (Figure S3B), and they form single-copy orthologs in all genomes except yeasts. Genes in Clade 1, including StuA homologs, are targets of the cyclic AMP (cAMP)-dependent protein kinase A (PKA) signal transduction pathway and were reported to be involved in dimorphic switch (Pan and Heitman 2000; Gimeno and Fink 1994), fungal spore development and the production of secondary metabolites (Lysøe et al. 2011).

Genes in Clade 2 and Clade 3 include *S. cerevisiae* Swi4 and Swi6, which form a protein complex and regulate genes essential during cell cycle progression from G1 to S phase (Koch et al. 1993), as well as meiosis (Son et al. 2016b). Genes in Clade 4 include homologs of *S. pombe* Bqt4 that connect telomeres to the nuclear envelope (Chikashige et al. 2009). Since this family of TFs is highly conserved across ascomycetes, similar functions can be proposed in *F. oxysporum*.

**DTT**, represented by the *S. cerevisiae* homolog Itc1, is a subunit of ATP-dependent Isw2p-Itc1p chromatin remodeling complex and is required for repression of early meiotic gene expression during the mitotic growth (Sugiyama and Nikawa 2001).

**RFX** represents a family of fungal TFs involved in DNA repair. Each strain encodes two orthologous copies, except *F. venenatum* encodes two copies within the RFX1 clade (Figure S3C). A major transcriptional repressor of DNA-damage-regulated genes in *S. cerevisiae*, Rfx1, is involved in DNA damage repair and replication checkpoint pathways (Lubelsky et al. 2005). In *F. graminearum*, Rfx1 is essential for maintaining the genome integrity (Min et al. 2014). The other copy, Rsc9 in *S. cerevisiae*, is a chromatin structure-remodeling complex RSC involved in transcription regulation and nucleosome positioning (Cairns et al. 1996; Hsu et al. 2003).

**NFYA** can bind to the CCAAT box. All strains maintain one copy of this family. The yeast homolog Hap2 induces the expression of mitochondrial electron transport genes (Olesen et al. 1991). *F. verticillioides* NFYA Hap2 is essential for fungal growth and the virulence on maize stalks (Ridenour and Bluhm 2014). The conservativeness suggests the functional importance of these TFs across *Ascomycota*, possibly linked to cellular machinery control (*e.g*. mitochondrial electron transport chain).

**MADS MEF2-like** family includes *S. cerevisiae* Rlm1, a component of the protein kinase C-mediated MAP kinase pathway involved in maintaining cell integrity (Jung et al. 2002). Rlm1 has a paralog from the whole genome duplication in *S. cerevisiae*, and all filamentous fungi encode one copy. In *F. verticilioides*, Mef2 plays a vital role in the sexual development (Ortiz and Shim 2013).

### 3. Minimal gene family contractions in FOSC partially caused by whole genome duplication in yeast

Four TF families, MATalpha_HMGbox, NOT4, MADS_SRF-like, and HSF (Heat Shock Factor), have an expansion score of less than 1, reflecting some level of gene family contraction among members of FOSC compared to the two yeast genomes (Figure S4).

**MATalpha_HMGbox** is a TF family that includes *S. cerevisiae* mating type protein alpha 1, a transcription activator that activates mating-type alpha-specific genes (Martin et al. 2010). All *F. oxysporum* Mat1-1 type strains contain this TF, but Mat1-2 strains do not. The contraction reflects the heterothallic mating strategy, even though sexual reproduction has not been observed in FOSC (Arie et al. 2000).

**NOT4** is a component of the multifunctional CCR4-NOT complex, a global transcriptional repressor of the RNA polymerase II transcription (Albert et al. 2002). This TF family remains a single copy in most genomes but is lost in some filamentous fungal genomes, including *A. nidulans*, *F. redolens*, *F. oxysporum* strains NRRL26365, MRL8666, and PHW726. It remains to be discovered why this gene is lost in some of these strains.

The contractions of the other two TF families, **MADS SRF-like** and **HSF**, are primarily caused by the whole genome duplication in yeast. In both cases, some degree of expansion was found in FOSC compared to other filamentous fungi (Figure S4).

**MADS SRF-like** is important for microconidium production and virulence in host plants, as reported in *M. oryzae* (Ding et al. 2020), and is essential for transcriptional regulation of growth-factor-inducible genes (Messenguy and Dubois 2003). The average copy number of phytopathogenic FOSC strains is 2.73, and the Fo5176 genome has the highest copy number of 6, while most other genomes only contain a single copy (Table S1).

**HSF** is a family of transcription factors that activate the production of many heat shock proteins that prevent or mitigate protein misfolding under abiotic/biotic stresses (Feder and Hofmann 1999). All non-FOSC filamentous fungi have three copies, while members of FOSC show expansion (*e.g*., Fo47: 4, Fol4287: 5, II5: 4, HDV274: 4, and Fo5176: 4) (Figure 3A-B). Interestingly, all expanded HSFs are phylogenetically close to Hsf1, which cluster together with the Hsf1 paralog of *Fusarium solani*, suggesting their horizontal transfer origin (Figure 3A). We then examined the *Hsf1* expression during the plant colonization (Guo et al. 2021). We found that the core copies of *Hsf1* of both strains Fo47 and Fo5176 were up-regulated during plant colonization. In contrast, the *Hsf1* accessory copies of these two strains were under opposite regulations, with Fo47 one being up-regulated and Fo5176 one being down-regulated post infection (Figure 3C), suggesting distinct regulatory adaptations after expansion. Here we noted that transcriptome data could be powerful in understanding the functional importance of TFs (see Section 6 for systematic analysis). In filamentous fungi, there are experimental reports for the other two clades. *Sfl1* is essential for vegetative growth, conidiation, sexual reproduction, and pathogenesis, as shown in *M. oryzae* (Li et al. 2011); *Skn7* is a regulator of the oxidative stress response and is essential for pathogenicity in *F. graminearum* (Jiang et al. 2015). Not surprisingly, both genes of Fo5176 and Fo47 were upregulated during plant colonization (Figure 3C).

**Figure 3.**
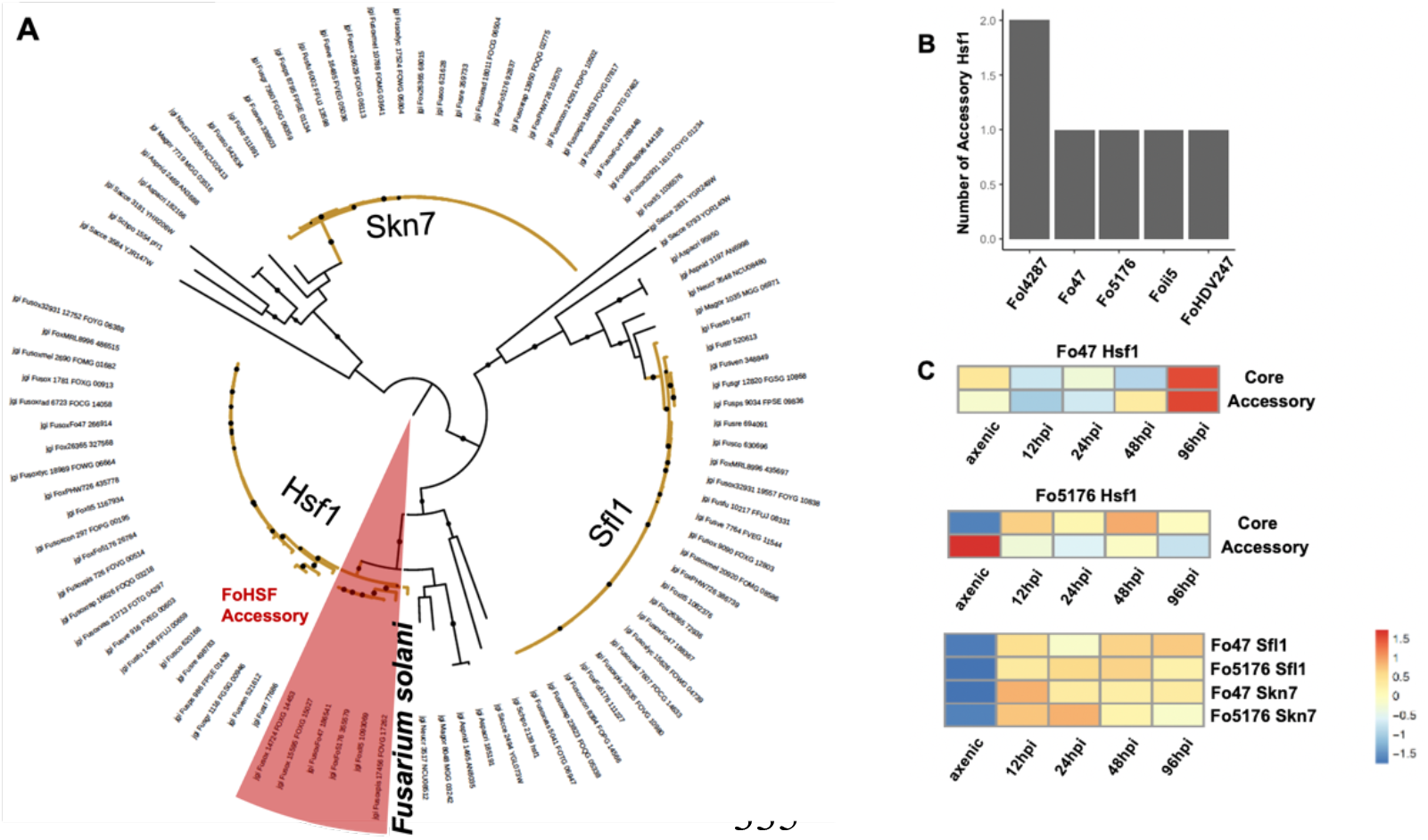
Evolutional trajectory of heat shock factors (HSFs) suggesting genome expansion and adaptation. (A) Phylograms of HSFs were constructed by maximum likelihood method with 1000 bootstraps. Branches of *Fusarium* HSFs were colored in yellow. Accessory HSFs of FOSC are shared in red. (B) Number of accessory HSFs in some FOSC genomes. (C) Expression of *HSF* genes during plant colonization (hpi indicates hours post inoculation), compared to axenic growth. Transcriptome data was previously described in Guo et al. 2021. See Materials and Methods for details of data reprocessing and visualization.

### 4. Significant TFome expansion in FOSC driven by a small number of exceedingly expanded TF families

#### 4.1. Gain-of-function among filamentous ascomycete fungi

Three TF families, CP2 (*EI_y_* = 2.73), HTH_AraC (*EI_y_* = 2), and HTH_Psq (*EI_y_* = 3.53), are absent in both yeast genomes, suggesting a gain of function among filamentous ascomycete fungi (Table S1). **CP2** has been studied in animal and fungal kingdoms with a function related to differentiation and development (Paré et al. 2012). Both **HTH_AraC** and **HTH_Psq** belong to helix-turn-helix (HTH) superfamily. First reported in bacteria, **HTH_AraC** is a positive regulator associated with the arabinose operon regulatory protein AraC (Schleif 2010; Gallegos et al. 1993; Bustos and Schleif 1993). **HTH_Psq**, as part of the eukaryotic Pipsqueak protein family, reported in vertebrates, insects, nematodes, and fungi, regulates the cell death (Siegmund and Lehmann 2002). Most FOSC genomes have a single copy of HTH_AraC, while the count of proteins containing the HTH_Psq ranges from 0 to 9 in the FOSC and ranges from 0 to 3 in other *Fusarium* relatives. Since the **HTH_Psq** domain also exists in transposases (Siegmund and Lehmann 2002), and ACs in FOSC are transposon-rich, it remains to be studied whether proteins containing the Psq domain are *bona fide* TFs.

#### 4.2. Seven exceedingly expanded TF families

Among others, seven TF families have expansion indexes greater than 2 (Table 2 and Figure 2B). Because of their drastic expansion, these seven families overall account for more than 75% of the total TFome. These families include Zn2-C6 (*EI_y_* = 15.09), bZIP (*EI_y_* = 5.80), and Znf_C2H2 (*EI_y_* = 4.15), Homeobox (*EI_y_* = 2.28), PAI2 (*EI_y_* = 3.42), NDT80 (*EI_y_* = 3.47), and bHLH (*EI_y_* = 3.48). All seven families show gradual expansion, reflected by the average copy number increment (FOSC > non-FOSC *Fusarium* > non-*Fusarium* filamentous fungi > yeasts, table S1). Furthermore, Zn2-C6 (44 in yeasts versus 671 in FOSC) and Znf_C2H2 (40 in yeasts versus 167 in FOSC) have the most drastic number increment along the evolutionary trajectory (Figure 2C and Table S1). Based on both high expansion index and large number increment, we considered Zn2-C6 and Znf_C2H2 as the most significantly expanded families.

The large copy number makes it hard to interpret functions from the protein domain annotation. Here we describe a couple of TFs reported in *F. oxysporum* and other systems and will introduce orthologous analysis to further survey the functionally validated TFs in the later section.

**Zn2-C6**, a fungal family TF (MacPherson et al. 2006), has the most significant expansion, reaching over 600 members among FOSC genomes and accounting for more than half of the total TFome. This group of TFs can form a homodimer and bind to the specific palindromic DNA sequence through direct contact with the major groove of the double-stranded DNA molecules (MacPherson et al. 2006). The versatility of this group of TFs can be achieved by domain shuffling and by changing the nucleotide binding specificity. In addition to the well-documented Ftf1 (Niño-Sánchez et al. 2016; van der Does et al. 2016; Ramos et al. 2007; Zuriegat et al. 2021; Zhao et al. 2020), five additional TFs within this family have been characterized in *F. oxysporum*, including Ctf1 (Rocha et al. 2008), Ctf2 (Rocha et al. 2008), Fow2 (Imazaki et al. 2007), XlnR (Calero-Nieto et al. 2007) and Ebr1 (Jonkers et al. 2014). They are involved in the development, metabolism, stress response, and pathogenicity.

**Znf_C2H2** is the most common DNA-binding motif found in the eukaryotic transcription factors (Fedotova et al. 2017). Five *F. oxysprum* TFs have been reported: *Czf1* (Yun et al. 2019), *Con7-1* (Ruiz-Roldán et al. 2015), *PacC* (Caracuel et al. 2003; Zhang et al. 2020), *ZafA* (López-Berges 2020) and *St12* (Asunción García-Sánchez et al. 2010; Rispail and Di Pietro 2009). Particularly, PacC was linked to the pathogenicity of both plant and human host (Zhang et al. 2020; Caracuel et al. 2003).

Other five families include **bZIP**, **Homeobox, PAI2, Ndt80** and **bHLH. bZIP** domain contains a region for sequence-specific DNA binding followed by a leucine zipper region required for dimerization (Bader and Vogt 2006). Three *F. oxysporum* bZIP TFs have been reported, including Atf1 (Li et al. 2013), Hapx (López-Berges et al. 2012), and MeaB (López-Berges et al. 2010), all of which are important for fungal pathogenicity. **Homeobox** is a DNA binding motif with a helix-turn-helix structure. In *S. pombe, Phx1* is a transcriptional coactivator that plays a role in yeast fission. In *M. oryzae, Hox* plays roles in the conidiation and appressorium development (Kim et al. 2009). **PAI2** is involved in the negative regulation of protease synthesis and sporulation of the *Bacillus subtilis* (Honjo et al. 1990). **Ndt80** is essential for completing meiosis in *S. cerevisiae* (Pierce et al. 2003; Tsuchiya et al. 2014) and *Ustilago maydis* (Doyle et al. 2016). It also promotes the expression of sporulation genes that are essential for the fulfillment of meiotic chromosome segregation (Hepworth et al. 1998). **bHLH** proteins form a large superfamily of transcriptional regulators found in almost all eukaryotes and function in critical developmental processes (Jones 2004). *F. graminearum* Gra2 is involved in the biosynthesis of phytotoxin gramillin (Bahadoor et al. 2018). *P. digitatum* encoding SreA is required for anti-fungal resistance and full virulence in citrus fruits (Liu et al. 2015).

#### 4.3. other families

Other 20 TF families (expanded but with *EI_y_* <= 2) account for 20% of the TFome; on average, each of these 20 families contains 9.6 copies in each genome examined (Table S1). These TFs are involved in chromatin remodeling and pheromone response, among other functions.

Four TF families are functionally linked to chromatin remodeling, including Bromodomain (*EI_y_* = 1.52), CBFA_NFYB (*EI_y_* = 1.35), Znf_RING-CH (*EI_y_* = 1.11), and ARID (*EI_y_* = 1.25). **Bromodomain** containing Spt7 is a crucial part of the SAGA complex in yeast. The SAGA complex is required to transcribe many genes in the genome. The bromodomain that is part of this subunit can recognize acetylated lysines of histones and eventually lead it to a more chromatin unwinding (Donczew et al. 2020). **CBFA_NFYB** is found in the proteins (*e.g*., *S. cerevisiae* Dls1) that regulate RNA polymerase II transcription through controlling chromatin accessibility (*e.g*., telomeric silencing) (Iida and Araki 2004). **Znf_RING-CH** has a functional connection to chromatin modification (*e.g*., *S. cerevisiae* Rkr1) (Braun et al. 2007). **ARID** is a 100 amino acid motif found in many eukaryotic TFs (Iwahara 2002). *S. cerevisiae* Swi1 plays a role in chromatin remodeling and is required to transcribe a diverse set of genes, including HO and Ty retrotransposons (Breeden and Nasmyth 1987; Hirschhorn et al. 1992).

**Ste12** is a family of TFs that regulate fungal development and pathogenicity (Rispail and Di Pietro 2010). These TFs are found only in the fungal kingdom. Ste12 binds to the DNA sequence that mediates pheromone response. It is involved in haploid mating and pseudohyphae formation in the diploid (Gancedo 2001). *F. oxysporum* Ste12 controls invasive growth and virulence downstream of the Fmk1-mediated MAPK cascade (Rispail and Di Pietro 2009). Except for *S. pombe* (missing one), every genome encodes one copy.

Among others, **Znf_NFX1** domain is found in the NK-X1, a repressor of the human disease-associated gene HLA-DRA (Song et al. 1994). **HMG_box** (high mobility group box) in *S. cerevisiae*, Spp41, is involved in negative expression regulation of spliceosome components (Maddock et al. 1994); Nhp6a is required for the fidelity of some tRNA genes (Braglia et al. 2007); Ixr1 is a transcriptional repressor that regulates hypoxic genes (Vizoso-Vázquez et al. 2012). One example of **Znf_GATA** is Fep1, a transcription factor that represses the expression of particular iron transporter genes under a high iron concentration (Kim et al. 2016). *S. cerevisiae* Mbf1, belonging to **Cro/C1-type HTH**, is a transcriptional coactivator (Takemaru et al. 1997).

### 5. Orthologous survey of TF families that were manually curated

To further understand expanded TFs and their impacts on transcriptional regulation, we curated a list of 102 TFs reported in literature focusing on *F. oxysporum*, *F. graminearum*, and other phytopathogenic fungi (Table S3 and examples as described in the previous section). Compared to this list of curated TFs using Orthofinder, we define 80 orthologous groups among *Fusarium* genomes (Table S4). 62 out of the 80 orthogroups have been identified using the above IPR-annotated pipeline, which enables the dissection of vastly expanded and high copy number TF families such as Zn2-C6 and Znf_C2H2, which are further mapped to 27 orthologous groups, including 17 in Zn2C6, 9 in Znf_C2H2, and 1 containing both Znf_C2H2 and Zn2-C6 domains (Table S4).

This effort also results in additional annotation to 18 TF families (Table S4), accounting for 32 genes per genome (3% of average *Fusarium* TFome size). These newly annotated TFs include homologs of those without domain annotation, *e.g*., disordered proteins *F. oxysporum* Ren1 (Ohara et al. 2004) and *M. oryzae* Som1 (Yan et al. 2011), and homologs of those with noncanonical TF domains such as **Ankyrin_rpt** and **WD40_repeat**.

We then directly compared *F. oxysporum* with its *Fusarium* relatives to calculate the expansion index as follows:

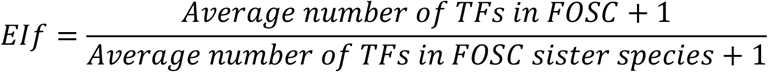

The *EI_f_* ranged, with the highest score being 3.54 (Fug, AreA_GATA) and the lowest being 0.5 (Fox1, Fork_head) (Table S4). Among these 80 orthogroups, 36 groups show high conservation (*EI_f_* = 1) as they are single-copy orthologs across *Fusarium*, among which ten were functionally validated in *F. oxysporum* (Table S4). 24 groups have gene contraction in *F. oxysporum* (*EI_f_* < 1). A total of 20 groups are expanded in *F. oxysporum* (*EI_f_* > 1, Table 3, Table S4), including five groups Fug1 (AreA_GATA, *EI_f_* = 3.54), Cos1 (Znf_C2H2, *EI_f_* = 2.8), Ftf1/Ftf2 (Zn2-C6, *EI_f_* = 2.7), Ebr1/Ebr2 (Zn2-C6, *EI_f_* = 2.5) and Ren1 (disordered, *EI_f_* = 2), with an *EI_f_* value equal or greater than 2. We also identified PacC (*EI_f_* = 1.57) as the second most expanded group within the highly expanded Znf_C2H2 family. We will further discuss these six groups (highlighted in bold, Table 3).

**Table 3.** Ortholog copy number and expansion index (*EI_f_*) of characterized and expanded TFs in *F. oxysporum*.

| TF | Reported species | References | Family | Overlap | Average_<br>Fo | Average_<br>non-Fo | El |
| --- | --- | --- | --- | --- | --- | --- | --- |
| **Ftf1/Ftf2** | ***F. oxysporum*** | **(Niño-Sánchez et al. 2016)** | **Zn2-C6** | **Yes** | **4.80** | **1.11** | **2.75** |
| **Ebr1/Ebr2** | ***F. oxysporum*** | **(Jonkers et al. 2014)** | **Zn2-C6** | **Yes** | **5.27** | **1.56** | **2.45** |
| Znf1 | *M. oryzae* | (Yue et al. 2016) | Zn2-C6 | Yes | 6.47 | 2.78 | 1.98 |
| Ctf2 | *F. oxysporum* | (Bravo-Ruiz et al. 2013) | Zn2-C6 | Yes | 2.93 | 1.33 | 1.69 |
| Fow2 | *F. oxysporum* | (Imazaki et al. 2007) | Zn2-C6 | Yes | 2.07 | 1.00 | 1.53 |
| Dep6 | *A. brassicicola* | (Wight et al. 2009) | Zn2-C6 | Yes | 0.93 | 0.67 | 1.16 |
| Pf2 | *A. brassicicola* | (Jones et al. 2019) | Zn2-C6 | Yes | 1.20 | 1.00 | 1.10 |
| Art1 | *F. verticillioides* | (Oh et al. 2016) | Zn2-C6 | Yes | 1.00 | 0.89 | 1.06 |
| Clta1 | *C. lindemuthianum* | (Dufresne et al. 2000) | Zn2-C6 | Yes | 1.07 | 1.00 | 1.03 |
| Fhs1 | *F. graminearum* | (Son et al. 2016a) | Zn2-C6 | Yes | 1.07 | 1.00 | 1.03 |
| **Cos1** | ***M. oryzae*** | **(Li et al. 2013)** | **Znf_C2H2** | **Yes** | **1.80** | **0.00** | **2.80** |
| **PacC** | ***F. oxysporum*** | **(Caracuel et al. 2003)** | **Znf_C2H2** | **Yes** | **2.13** | **1.00** | **1.57** |
| **Fug1** | ***F. verticillioides*** | **(Ridenour and Bluhm 2017)** | **AreA_GATA** | **No** | **7.27** | **1.33** | **3.54** |
| **Ren1** | ***F. oxysporum*** | **(Ohara et al. 2004)** | **disordered** | **No** | **3.00** | **1.00** | **2.00** |
| Tri10 | *F. graminearum* | (Jiang et al. 2016) | Fun_TF | No | 1.13 | 0.33 | 1.60 |
| Ltf1 | *B. cinerea* | (Schumacher et al. 2014) | Znf_GATA | Yes | 4.00 | 2.44 | 1.45 |
| Ndt80 | *U. maydis* | (Doyle et al. 2016) | NDT80 | Yes | 1.73 | 1.11 | 1.29 |
| Hap3p | *F. verticillioides* | (Ridenour and Bluhm 2014) | CBFA_NFYB | Yes | 1.33 | 1.00 | 1.17 |
| Sod1 | *F. oxysporum* | (Wang et al. 2021) | SOD_Cu_Zn | No | 1.47 | 1.22 | 1.11 |
| Prf1 | *F. oxysporum* | (Mendoza-Mendoza et al. 2009) | HMG_box | Yes | 1.07 | 1.00 | 1.03 |

Both **Ftf1/Ftf2** and **Ebr1/Ebr2** belong to the Zn2-C6 family and contribute directly to the fungal virulence (Michielse et al. 2009; van der Does et al. 2016; Ramos et al. 2007). Deletion of accessory copy Ftf1 reduced the pathogenicity of *F. oxysporum* f. sp. *phaseoli* (Ramos et al. 2007), highlighting the direct functional involvement of AC TF in virulence. In Fol, deletion of either Ftf1 (AC encoding) or Ftf2 (CC encoding) reduced the virulence towards the host (de Vega-Bartol et al. 2011; Niño-Sánchez et al. 2016). Constitutive expression of either *Ftf1 or Ftf2* induced the expression of effector genes (van der Does et al. 2016). The core copy Ftf2 is conserved among all *Fusarium* species, and the AC copy Ftf1 is only found in *F. oxysporum* and *Fusarium redolens* (Figure 4). **Ebr1** and paralogues are responsible for virulence and general metabolism. In *F. oxysporum*, Ebr1 is found as multiple homologs, whereas in *F. graminearum*, it is seen as a single copy (Jonkers et al. 2014). In *F. oxysporum*, three paralogous copies, Ebr2, Ebr3, and Ebr4, are encoded in ACs and regulated by core copy Ebr1. The importance of the core paralog has been shown by the reduced pathogenicity and growth defects when it was knocked out (Jonkers et al. 2014). It is worth noting that the *Ebr2* coding sequence driven by an *Ebr1* promoter was able to rescue the *Ebr1* knockout mutation, indicating some functional redundancy of this family.

**Figure 4.**
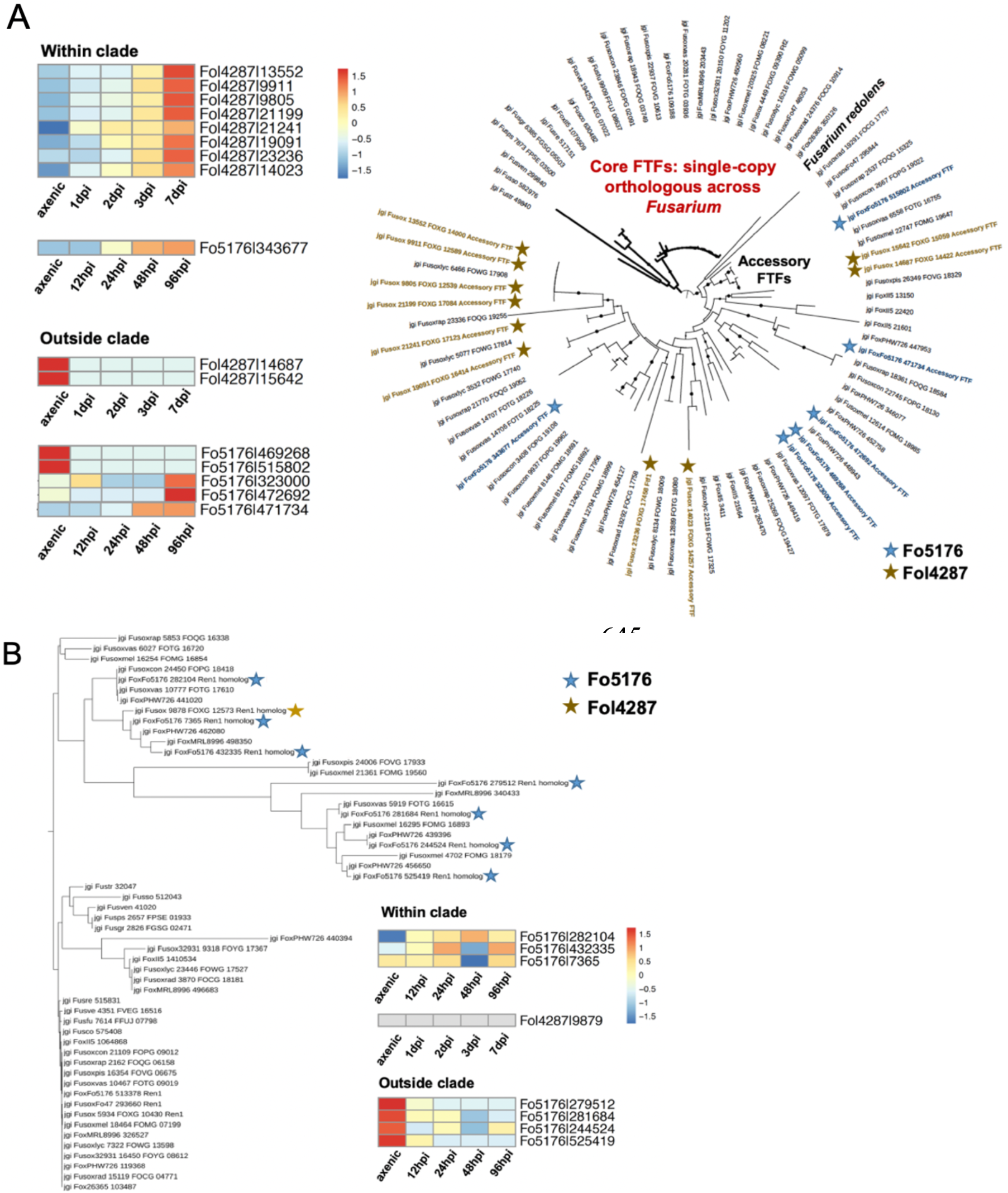
Unique expansion of some TFs, driven by ACs, may provide clues to host-specific adaptation. RNA-seq data were previously described (Guo et al., 2021; Redkar et al., 2021). (A) Ftf1, the TF involved in the tomato pathogenicity is most significantly expanded (10 copies of accessory FTFs) in the tomato pathogen Fol4287 genome and the expression of eight out of 10 were induced during plant colonization. (B) Ren1 is most significantly expanded (seven copies of accessory RENs) in the Arabidopsis pathogen Fo5176 genome, and two of them were induced during plant colonization.

Both **Cos1** and **PacC** belong to the Znf_C2H2 family. Mutation to *M. oryzae Cos1* resulted in developmental failure of the conidiophores (Li et al. 2013). Furthermore, mutation to *Cos1* aggravated the plant infection of leaf blades and sheaths, indicating a negative role in the pathogenicity (Zhou et al. 2009). PacC is an important pH-responsive TF in *F. oxysporum* (Caracuel et al. 2003; Zhang et al. 2020). PacC homologs are expanded in clinical strains (average accessory copy number 3.7) of FOSC, compared to non-clinical strains (average accessory copy number 0.5), while all *Fusarium* relatives’ genomes examined only contain a single copy of core PacC. Our previous study revealed that in *F. oxysporum* clinical strains, the expression of one expanded *PacC* gene on ACs was induced and the protein localized in the nucleus at mammalian physiological pH (7.4), indicating a potential role in host adaptation (Zhang et al. 2020). Interestingly, the induction of AC-encoding *PacC* genes was CC-encoding *PacC* gene-dependent, as the induction disappeared in the CC-encoding *PacC* knockout mutant, further supporting a cross-talking between core and accessory TFs (Yang 2020). Similar to *EBR1*, the expression of AC *PacC* genes is much lower than that of the CC *PacC* gene, and knockouts of one AC *PacC* gene affected a small subset of genes compared with the CC *PacC* knockout, which has a broader effect on cellular processes (Yang 2020).

**Fug1** has a role in pathogenicity (maize kernel colonization) and fumonisin biosynthesis in *F. verticillioides* (Ridenour and Bluhm 2017). In addition, the deletion of *Fug1* increased sensitivity to the antimicrobial compound 2-benzoxazolinone and to hydrogen peroxide, which indicates that Fug1 plays a role in mitigating stresses associated with the host defense (Ridenour and Bluhm 2017). Neither core copies nor accessory copies of these two genes were experimentally examined in FOSC. **Ren1** is a disordered protein with no IPR functional domain. The expansion score *EI_f_* = 2 suggests a unique expansion among FOSC. However, the only reported study on its function is in *F. oxysporum f. sp. melonis* regulating the development of the conidiation (Ohara et al. 2004).

### 6. Transcriptome analysis to probe the essential TFs during host colonization

To understand the functional importance of FOSC TFs, we take advantage of two recently reported transcriptomics datasets (Redkar et al. 2022; Guo et al. 2021), including pathogenic interactions (Fo5176 infecting Arabidopsis and Fol4287 infecting tomato) and endophytic interactions (Fo47 colonizing Arabidopsis) (Supplemental Dataset).

We first asked what proportion of genes was expressed in conserved and expanded categories (Table S5). We found that almost all genes (58 out of 60) within the conserved category (Group 1) were consistently expressed (TPM > 1 across all conditions), supporting their general roles in controlling life processes. Within the expanded category (Group 3), the proportion of genes being consistently expressed ranges from 41% to 59% for core TFs, and ranges from 5% to 16% for accessory TFs. With a less strict filter (TPM > 1 at minimum 1 condition), we found that all genes within the conserved category were expressed. Within the expanded category, the proportion of genes being expressed accounts for 93% of core TFs across all strains and ranges from 49% to 67% for accessory TFs. When we compared genes being consistently expressed versus genes being expressed at a minimum one condition, the more dramatic number increase for the expanded category (especially when we only consider the accessory TFs) highlighted that the expanded category, especially the accessory TFs, are more likely to be conditional expressed, further supporting their role in niche adaptation.

With the goal of examining the expression and probing important core and accessory TFs, we aimed to develop filtering parameters. Since most validated TFs were reported in the reference Fol4287 strain, we first reviewed for the reported TFs, both core, and accessory, the expression pattern during Fol4287 infecting tomatoes (Table S6). Out of 27 TFs encoded on the core genome, 18 show up-regulation (defined by at least three out of four *in planta* conditions show up-regulation compared to the axenic growth) during plant colonization from 1 day post-inoculation (dpi) to 7 dpi, consistent with their reported roles in pathogenicity. The accessory *Ftf1* has been exclusively demonstrated to play essential functions in fungal pathogenicity (Niño-Sánchez et al. 2016). Eight of 10 accessory *Ftf*s were upregulated using the same criteria during plant colonization. Our results illustrate the power of using transcriptome data to probe the functionally important players during plant colonization/infection.

We further developed strict criteria to filter important TFs from TFome (Figure S5), by which half (nine) of previously described upregulated core TFs meet the ‘core’ criteria, and all eight up-regulated accessory *Ftf*s meet the ‘accessory’ criteria (Table S6). We then apply such measures to the transcriptome of all the TFomes of Fol4287, Fo5176, and Fo47 to probe two types of TFs: 1) The conserved core TFs related to plant colonization; 2) Expanded accessory TFs related to host-specific pathogenicity.

Fol4287, Fo5176, and Fo47 upregulated 95, 62, and 44 core TFs during plant colonization. Among them, ten copies are highly conserved (Table S7), as they are single-copy orthologs across all 15 *F. oxysporum* strains. Two out of ten were previously reported, *Fow2* and *Sfl1*. Fow2, Zn2C6 TF, is required for full virulence but not hyphal growth and conidiation in *F. oxysporum* f. sp. *melonis* (Imazaki et al. 2007). The downstream targets of Fow2 remain unknown in *F. oxysporum*, thus meriting further analysis. Sfl1 has been described in the previous section and is essential for vegetative growth, conidiation, sexual reproduction, and pathogenesis, as shown in *M. oryzae* (Li et al. 2011). The functions of FoSfl1 remain to be validated.

Fol4287, Fo5176, and Fo47 upregulated 29, 34, and 9 accessory TFs. *Ftf1* and *Ren1* are particularly interesting (Figure 4 and Table S8). Though Ftfs have been shown to play an essential role in pathogenicity in Fol, whether this pathway is restricted to the same strain remains a question. Compared to Fol4287 which contains ten accessory *Ftf*s and eight were upregulated during plant colonization, Fo5176 includes six copies of accessory *Ftf*s, but only one copy was upregulated. Interestingly, eight upregulated Fol4287 and one upregulated Fo5176 *Ftf*s are clustered together (Figure 4). The unique expansion with regulatory adaptation (*i.e*., fine-tuned expression regulation) seems to be restricted to Fol4287 but not another pathogenic strain, Fo5176, when they infect the hosts. Among Fo5176 expanded TFs, we identified *Ren1*. Compared to Fol4287 which encodes only one accessory *Ren1* that was not upregulated during plant colonization, Fo5176 encodes seven accessory copies, among which two were upregulated (Figure 4). Though functional validation is needed, the strain-specific expansion followed by fine-tuned expression regulation when infecting host species exists and likely contributes to the host-specific pathogenicity.

## DISCUSSION

For a soilborne pathogen with strong host specificity like FOSC, the adjustment of growth and cell cycle control in response to environmental cues is likely essential for survival. At the same time, expanded families likely contribute to the enhanced functions related to niche adaptation. TFs transmit external and internal signals and regulate complex cellular signaling responses to the sensed stimuli. Transmitted through the soil and vascular wounds of plants causing vascular wilt (Gordon 2017), *F. oxysporum* must adapt to stresses encountered both outside and inside its host. Therefore, it is not surprising to see that genomes of FOSC have larger TFome than other fungi included in the study. The expansion of TFs among FOSC resulted in a positive correlation between the total number of proteins and the size of the fungal TFome, which was also observed before (Shelest 2017).

A total of 14 TF families that control the global transcriptional event, such as TBP, are highly conserved within the ascomycete fungal lineages. Conserved regulatory mechanisms revealed through this study suggest that the plant colonization process could be a common process among FOSC strains regardless of their host-specific pathogenesis. The notion was also supported by recent studies that highlighted the ability of FOSC as a root colonizer facilitated by the conserved genomics components (Martínez-Soto et al. 2022; Redkar et al. 2022).

In contrast to these stable TFs, 30 families are expanded in various degrees and most significant expansions occurred in Zn2-C6 and Znf_C2H2 TF families among FOSC genomes. The number of Zn2-C6 TFs increases significantly (with the highest expansion score) and makes up most of the TFs (56.7%) found within the FOSC TFome. For example, *Ftf1*, a TF belonging to the Zn2-C6 and involved in the tomato pathogenicity, is most significantly expanded (10 copies of accessory *Ftf*s) in the tomato pathogen Fol4287 genome, and the expression of eight out of 10 was induced during plant colonization. The continuous expansion suggests the functional importance of these understudied TFs, further supported by the genetic studies (Niño-Sánchez et al. 2016; van der Does et al. 2016; Ramos et al. 2007; Zuriegat et al. 2021; Zhao et al. 2020) and their induction during host invasion revealed by our RNA-seq data.

Unique expansion of some TFs, driven by ACs, may provide a clue to host-specific interactions. Acquiring additional TFs will modify existing regulatory pathways. No question, this will require the fine-tuning of existing networks for this group of organisms to successfully adapt to different hosts under diverse environments. A previous survey of kinome (the complete set of protein kinases encoded in an organism’s genome) among FOSC and other Ascomycetes revealed a positive correlation between the size of the kinome and the size of the genome (DeIulio et al. 2018), exactly the same as we reported here for TFomes. As kinases and TFs are key regulators that modulate all important signaling pathways and are essential for the proper functions of almost all molecular and cellular processes. Strong correlations among kinome and TFome suggest an ordered, instead of chaotic, recruitment and establishment of ACs among FOSC genomes.

This realization further emphasizes the importance of additional functional studies. Reverse genetics is a powerful tool in defining the functional importance of a TF. For example, TF Ren1, a disordered protein, was identified by genetic and molecular characterization (Ohara et al. 2004). This TF is most significantly expanded (seven copies of accessory *Ren*s) in the Arabidopsis pathogen Fo5176 genome, and two of them were induced during plant colonization. Experiments such as chromatin immunoprecipitation sequencing (CHIP-Seq) and DNA affinity purification sequencing (DAP-seq) to profile the cis-regulatory elements globally are high throughput approaches to define specific binding sites (cis-regulatory elements) of TFs. DAP-seq was used successfully to profile the Cistrome for the entire TFome of the bacterial organism (Baumgart et al. 2021), holding the promise for a better understanding of transcriptional regulation in the fungal model *F. oxysporum*. TFs can function individually or with other proteins in a complex, and can act as an activator that promotes transcription or a repressor that blocks the recruitment of RNA polymerase. Therefore, defining specific functions of these identified binding sites through DAP-seq can be difficult. Gene regulatory networks based on gene co-expression and other phenotypic and multi-omics data as reported in *Fusarium* (Guo et al. 2016, 2020) can add more resolution to these complex regulatory processes. However, the ultimate understanding of the regulatory roles of each TF will come from careful molecular and biochemical characterization.

A systematic understanding of transcriptional regulation is essential to get the fine-tuned footprint of the gene regulatory network. Our study not only offered a comprehensive look at the regulation from the evolutionary perspective, but also provided an easily implemented computational pipeline to compare TFs and other functional groups in fungi. A better understanding of their functions would not only inform *Fusarium* biology but also could be extrapolated to other filamentous fungi and complex basidiomycetes.

## SUPPLEMENTAL MATERIALS

Table S1. TF-type DNA-binding domains used to filter TFome

Table S2. TFome annotations across 30 genomes

Table S3. Curated TFs in phytopathogens

Table S4. Ortholog copy number of characterized TFs of phytopathogens across 30 genomes

Table S5. Number of TFs being expressed

Table S6. Orthologs of reported TFs in Fol4287

Table S7. Highly conserved TFs that are constantly upregulated during plant colonization

Table S8. Strain-specific accessory TFs that are upregulated during plant colonization

Figure S1. TFome annotation pipelines

Figure S2. ACs contribute to the FOSC TFome expansion

Figure S3. Phylograms of three conserved families

Figure S4. Minimal gene family contractions in FOSC partially caused by whole genome duplication in yeast

Figure S5. The pipelines to probe the functional important TFs by RNA-seq data

Supplemental Dataset. Normalized read counts of RNA-seq datasets

## ACKNOWLEDGMENTS

We thank the Massachusetts Green High-Performance Computing Center for providing high-performance computing capacity.

## AUTHOR CONTRIBUTIONS

Conceptualization, H.Yu and L-J.M.; Methodology, Analysis and Interpretation, all authors; Resources, S.H., R.H. and I.G.; Writing – H.Yu and L-J.M. with input from all authors; Supervision, L-J.M.; Funding Acquisition, L-J.M.

