## Supplementary material for "Conservation and Expansion of Transcriptional Factor Repertoire in the *Fusarium oxysporum* Species Complex": TFome_Supplemental Figures.pptx

### Slide 1
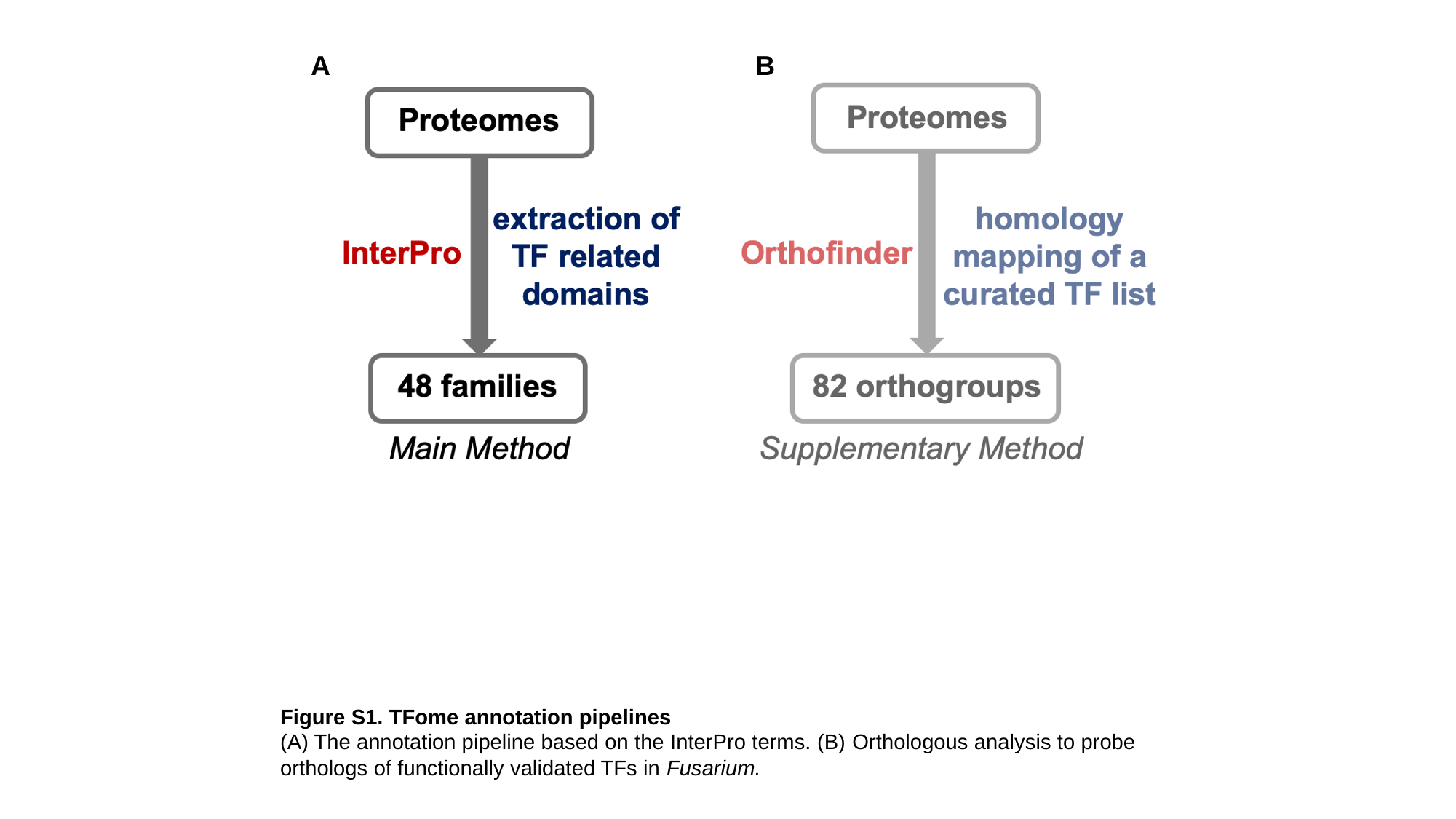

A
B
Figure S1. TFome annotation pipelines
(A) The annotation pipeline based on the InterPro terms. (B) Orthologous analysis to probe orthologs of functionally validated TFs in Fusarium.

### Slide 2
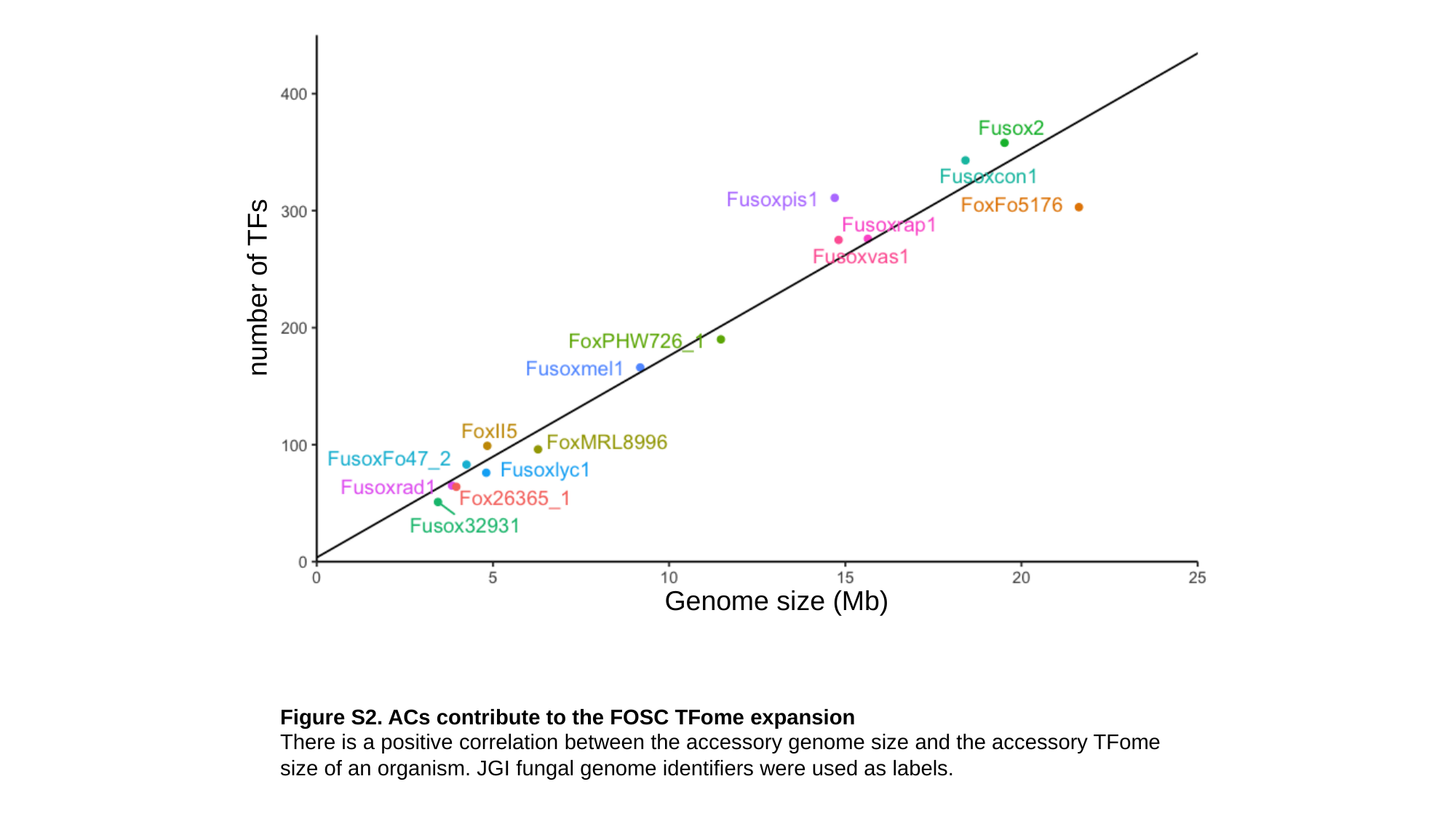

number of TFs
Genome size (Mb)
Figure S2. ACs contribute to the FOSC TFome expansion
There is a positive correlation between the accessory genome size and the accessory TFome size of an organism. JGI fungal genome identifiers were used as labels.

### Slide 3
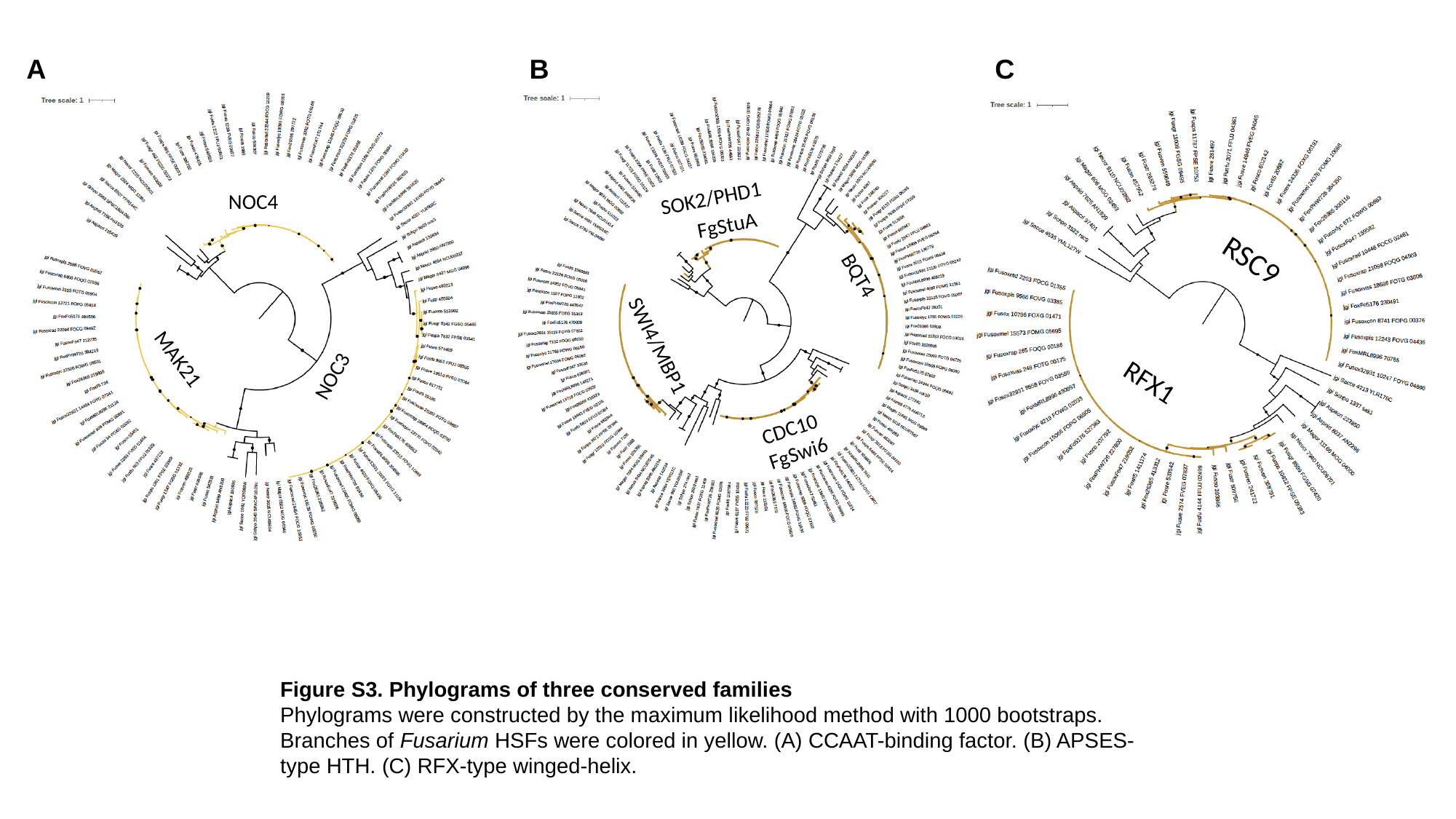

C
A
B
SOK2/PHD1
NOC4
FgStuA
RSC9
BQT4
NOC3
SWI4/MBP1
RFX1
MAK21
CDC10
FgSwi6
Figure S3. Phylograms of three conserved families
Phylograms were constructed by the maximum likelihood method with 1000 bootstraps. Branches of Fusarium HSFs were colored in yellow. (A) CCAAT-binding factor. (B) APSES-type HTH. (C) RFX-type winged-helix.

### Slide 4
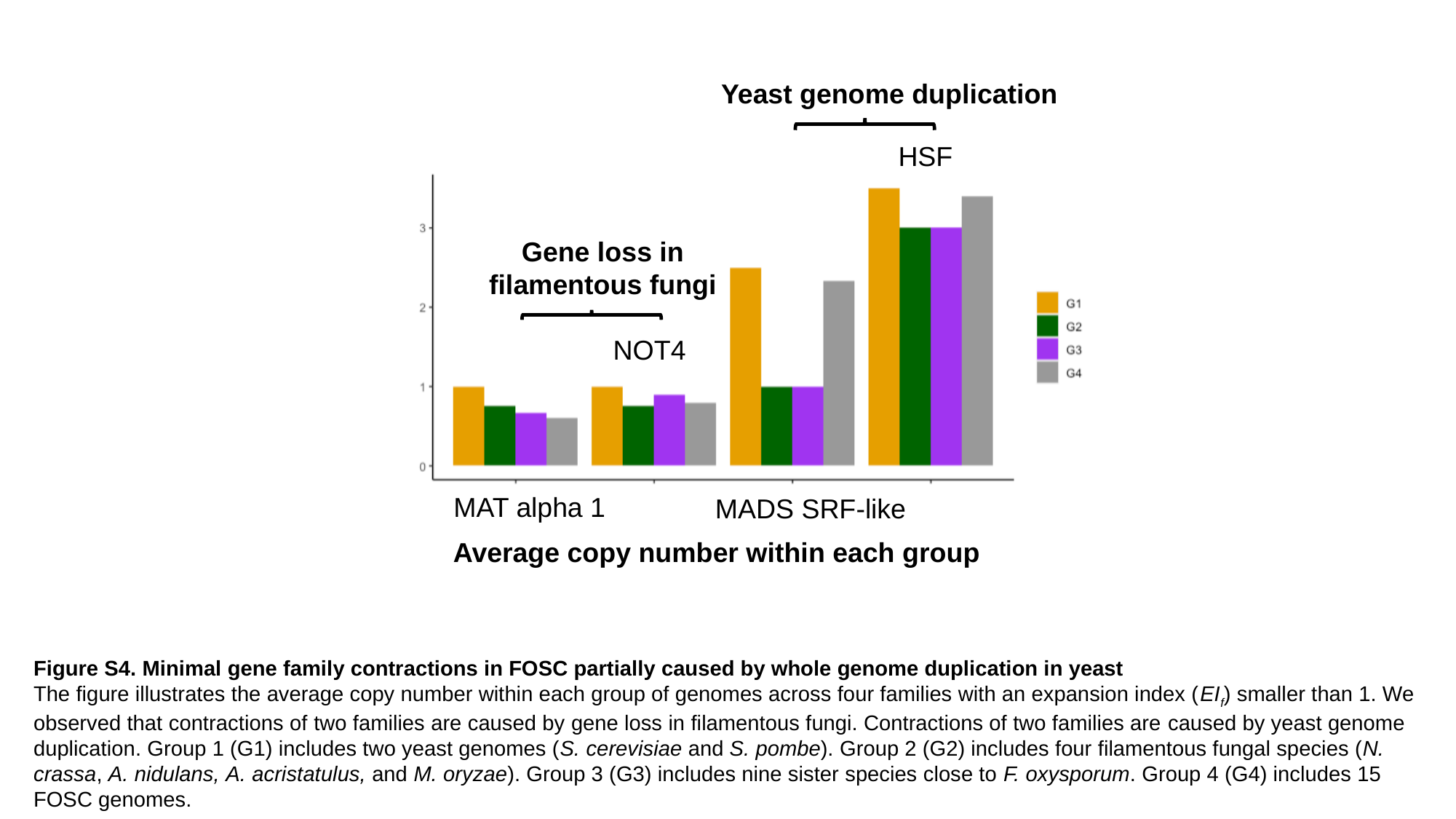

Yeast genome duplication
HSF
Gene loss in filamentous fungi
NOT4
MAT alpha 1
MADS SRF-like
Average copy number within each group
Figure S4. Minimal gene family contractions in FOSC partially caused by whole genome duplication in yeast
The figure illustrates the average copy number within each group of genomes across four families with an expansion index (EI_f_) smaller than 1. We observed that contractions of two families are caused by gene loss in filamentous fungi. Contractions of two families are caused by yeast genome duplication. Group 1 (G1) includes two yeast genomes (S. cerevisiae and S. pombe). Group 2 (G2) includes four filamentous fungal species (N. crassa, A. nidulans, A. acristatulus, and M. oryzae). Group 3 (G3) includes nine sister species close to F. oxysporum. Group 4 (G4) includes 15 FOSC genomes.

### Slide 5
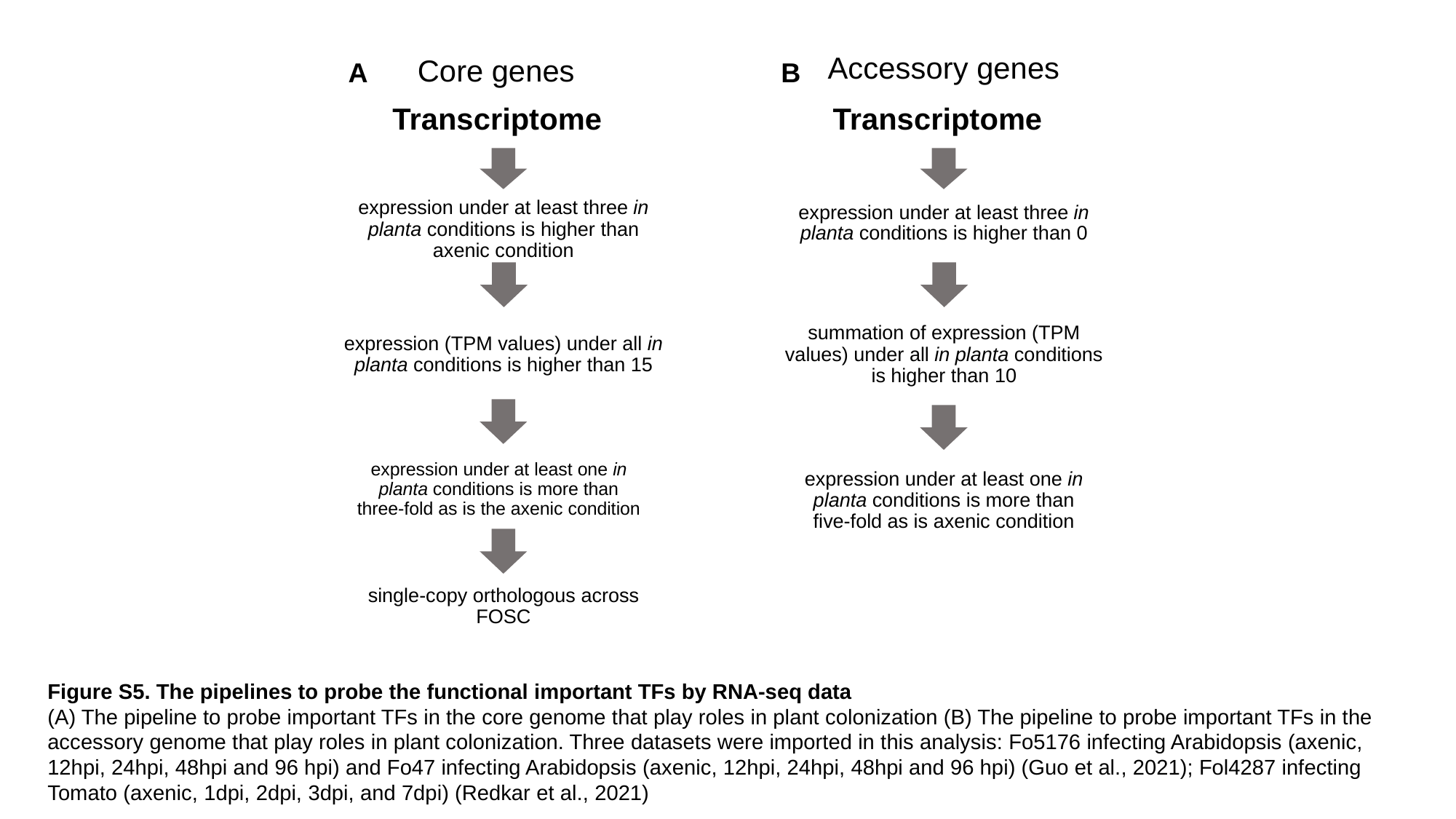

Accessory genes
Core genes
A
B
Transcriptome
Transcriptome
expression under at least three in planta conditions is higher than axenic condition
expression under at least three in planta conditions is higher than 0
expression (TPM values) under all in planta conditions is higher than 15
summation of expression (TPM values) under all in planta conditions is higher than 10
expression under at least one in planta conditions is more than three-fold as is the axenic condition
expression under at least one in planta conditions is more than five-fold as is axenic condition
single-copy orthologous across FOSC
Figure S5. The pipelines to probe the functional important TFs by RNA-seq data
(A) The pipeline to probe important TFs in the core genome that play roles in plant colonization (B) The pipeline to probe important TFs in the accessory genome that play roles in plant colonization. Three datasets were imported in this analysis: Fo5176 infecting Arabidopsis (axenic, 12hpi, 24hpi, 48hpi and 96 hpi) and Fo47 infecting Arabidopsis (axenic, 12hpi, 24hpi, 48hpi and 96 hpi) (Guo et al., 2021); Fol4287 infecting Tomato (axenic, 1dpi, 2dpi, 3dpi, and 7dpi) (Redkar et al., 2021)
